# The Arabidopsis SWEET1 and SWEET2 uniporters recognize similar substrates despite differences in subcellular localization

**DOI:** 10.1101/2023.05.30.542931

**Authors:** Sojeong Gwon, Jihyun Park, AKM Mahmudul Huque, Lily S. Cheung

## Abstract

Sugars Will Eventually be Exported Transporters (SWEETs) are central for sugar allocation in plants. The SWEET family is vast, with approximately 20 homologs in most plant genomes. Despite extensive research on their structures and molecular functions, it is still unclear how diverse SWEETs recognize their substrates. Previous work using SweetTrac1, a biosensor constructed by the intramolecular fusion of a conformation-sensitive fluorescent protein in the plasma membrane transporter SWEET1 from *Arabidopsis thaliana*, identified common features in the transporter’s substrates. Here, we report SweetTrac2, a new biosensor based on the Arabidopsis vacuole membrane transporter SWEET2 and use it to explore the substrate specificity of this second protein. Our results show that SWEET1 and SWEET2 recognize similar substrates but some with different affinities. Sequence comparison and mutagenesis analysis support the conclusion that the differences in affinity depend on non-specific interactions involving key residues in the binding pocket. Furthermore, SweetTrac2 can be an effective tool for monitoring sugar transport at vacuolar membranes that would be otherwise challenging to study.

## Introduction

Sugars are undoubtedly the main carbon source in plants. Efficiently mobilizing sugars from photosynthetic organelles to other organs and tissues is crucial for many biological processes. Members of the Sugar Will Eventually be Exported Transporters (SWEET) family are uniporters found in both plasma and vacuolar membranes that play crucial roles in plant growth and development, pathogen susceptibility, and stress tolerance (1,2). Extensive research is being conducted as a result to capitalize on these proteins for crop improvement.

SWEETs are also small, making them good molecular models for exploring sugar recognition mechanisms. Most eukaryotic SWEETs have seven transmembrane domains, while the bacterial homologs, the SemiSWEETs, have only three transmembrane domains and form dimers to complete the sugar translocation path (3-5). SWEETs are smaller than other sugar transporters with available crystal structures, such as the human GLUT1, the *Escherichia coli* LacY, and the *Vibrio parahaemolyticus* SGLT (6,7). The minimal size facilitates cloning, purification, heterologous expression, and mutagenesis studies. In fact, mutations of 13% of the amino acids in the Arabidopsis SWEET1 (AtSWEET1) protein have been characterized so far (2), a large fraction compared with most other transporters.

However, unlike plasma membrane transporters like AtSWEET1, vacuolar membrane transporters are considerably more difficult to characterize. Most studies on vacuolar proteins start with the isolation of vacuoles, which are very fragile and require careful handling (8,9). Purifying and testing the transporters afterwards is even more challenging since they only represent about 1% of the total cell protein and must go through labor-intensive solubilization and reconstitution steps (10).

When sufficient vacuolar sugar transporters are successfully incorporated into vesicles, radiolabeled substrates are commonly used to test uptake and efflux. If sugar translocation is accompanied by the movement of charged ions (e.g., H^+^), sophisticated methods such as solid-supported membrane (SSM)-based electrophysiology could be used (11). Despite the complications associated with studying vacuolar sugar transporters, they should not be neglected, as the vacuole is the main sugar storage organelle and regulates sugar dynamics in the cell (12).

Transporter biosensors are chimeras of transporters and fluorescent proteins that translate the conformational changes of the transporter into a fluorescence response, offering an alternative to traditional characterization methods (13). These biosensors can be a powerful tool to study transporter activity in real-time, and a few have been successfully engineered for plant proteins, including SweetTrac1, a biosensor based on AtSWEET1 (14-16). We previously proposed a kinetic model that describes the dynamic response of SweetTrac1 to D-glucose (14). More recently, we employed SweetTrac1 and cheminformatics to investigate the substrate specificity of AtSWEET1, enabling us to find potential substrates of AtSWEET1 in a quick and easy manner without the need for radiolabeled chemicals or individual intracellular sensors for each chemical (17).

In this work, we convert the vacuole membrane localized AtSWEET2 uniporter into the biosensor SweetTrac2 and characterize its substrate specificity. We show that despite their distinct subcellular localization, AtSWEET1 and AtSWEET2 recognize similar substrates, albeit with different affinities. Furthermore, sequence comparison of AtSWEET1 and AtSWEET2 and mutagenesis analysis identified three residues responsible for the differences in affinities.

## Results

### Generating and photophysically characterizing SweetTrac2

AtSWEET2 is a vacuole transporter that facilitates sugar storage in roots (18). AtSWEET1 and AtSWEET2 are both classified as clade I SWEETs that prefer hexoses as substrates (19). Given the 44% sequence identity of these two proteins (Fig. 1A), we hypothesized that transferring the circularly permutated, superfolded green fluorescent protein (cpsfGFP) and linkers of SweetTrac1 into the same position in AtSWEET2 would result in another successful biosensor. As predicted, this approach generated the new biosensor SweetTrac2, which responded to the addition of glucose in a similar manner to SweetTrac1 (Fig. 1B).

**Figure 1.**
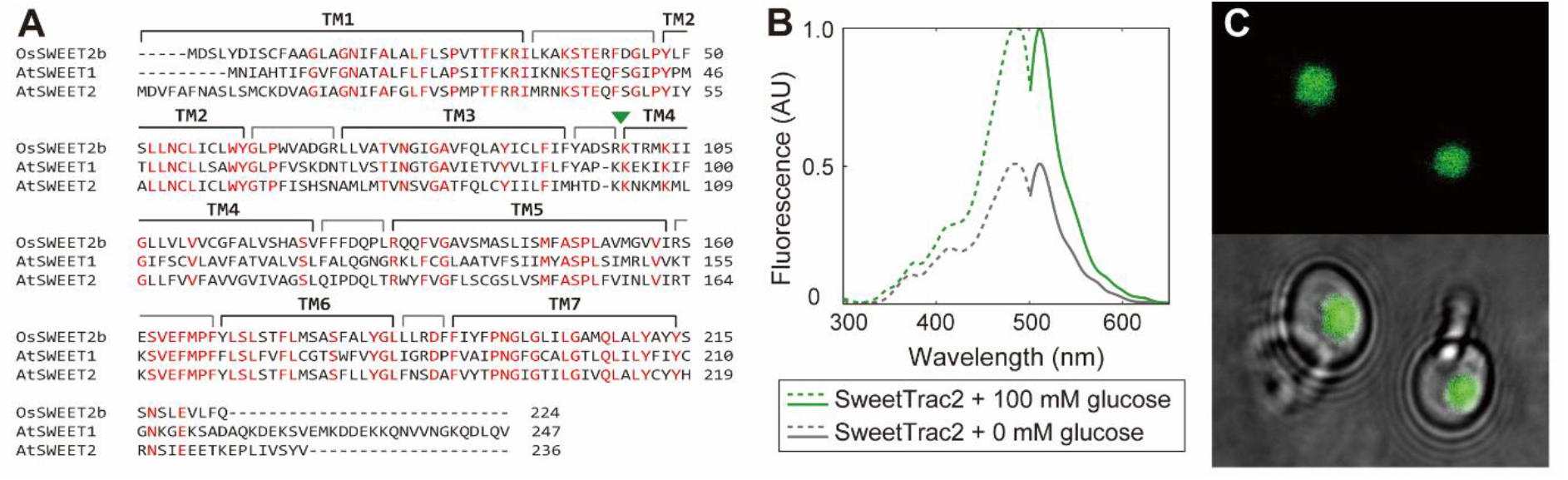
Characterization of the SweetTrac2 biosensor. (A) Multiple sequence alignment of OsSWEET2b, AtSWEET1, and AtSWEET2. Highlighted in red are amino acids that are conserved across the three proteins. Transmembrane domains (TM) based on an alignment with OsSWEET2b are marked above the sequence. Green arrowhead indicates the positions where the linkers and cpsfGFPs were inserted for the construction of SweetTrac1 and SweetTrac2. (B) Normalized fluorescence excitation and emission spectra of SweetTrac2 (455 nm excitation, 530 nm emission). Dashed lines illustrate excitation, and solid lines illustrate emission. (C) Localization of SweetTrac2 to the vacuolar membrane in yeast cells.

Subcellular localization of both c-terminus tagged AtSWEET1 and AtSWEET2 in yeast cells mirrors that of the natural transporters *in planta* (18,20). Similarly, SweetTrac1 localized to the plasma membrane (14), while SweetTrac2 localized to the vacuole (Fig. 1C).

Spectra analysis of SweetTrac2 revealed two excitation maxima—a major peak from the deprotonated chromophore at a wavelength of ∼490 nm and a minor peak from the protonated chromophore at a wavelength of ∼410 nm. A single emission maximum was observed at a wavelength of ∼515 nm (Fig. 1B). The peak fluorescence intensity increases with glucose addition, and no shift in excitation and emission maxima was observed (Fig. 1B).

### Characterizing the substrate specificity of AtSWEET2

In our previous work, we expressed SweetTrac1 in yeast, screened a custom library of 162 sugar and sugar analogs, and performed a cheminformatics analysis of the results. We identified 15 chemicals capable of inducing a fluorescence response, suggesting they can bind the transporter’s substrate binding pocket (Fig. 2). Three of these hits (D-glucose, D-fructose, and D-mannose) were known substrates of AtSWEET1, and we confirmed their cellular uptake using radiolabeled versions of these sugars. We also confirmed that AtSWEET1 could mediate the cellular uptake of three other hits (1-deoxynojirimycin, voglibose, and 1-thio-D-glucose), which have adverse effect on yeast growth (17). Unfortunately, this previous study did not clarify whether the remaining nine hits were substrates of AtSWEET1 or competitive inhibitors (capable of binding the transporter but not translocated to the cytosolic side of the membrane), as radiolabeled versions of them are not commercially available and they did not hinder cell growth.

**Figure 2.**
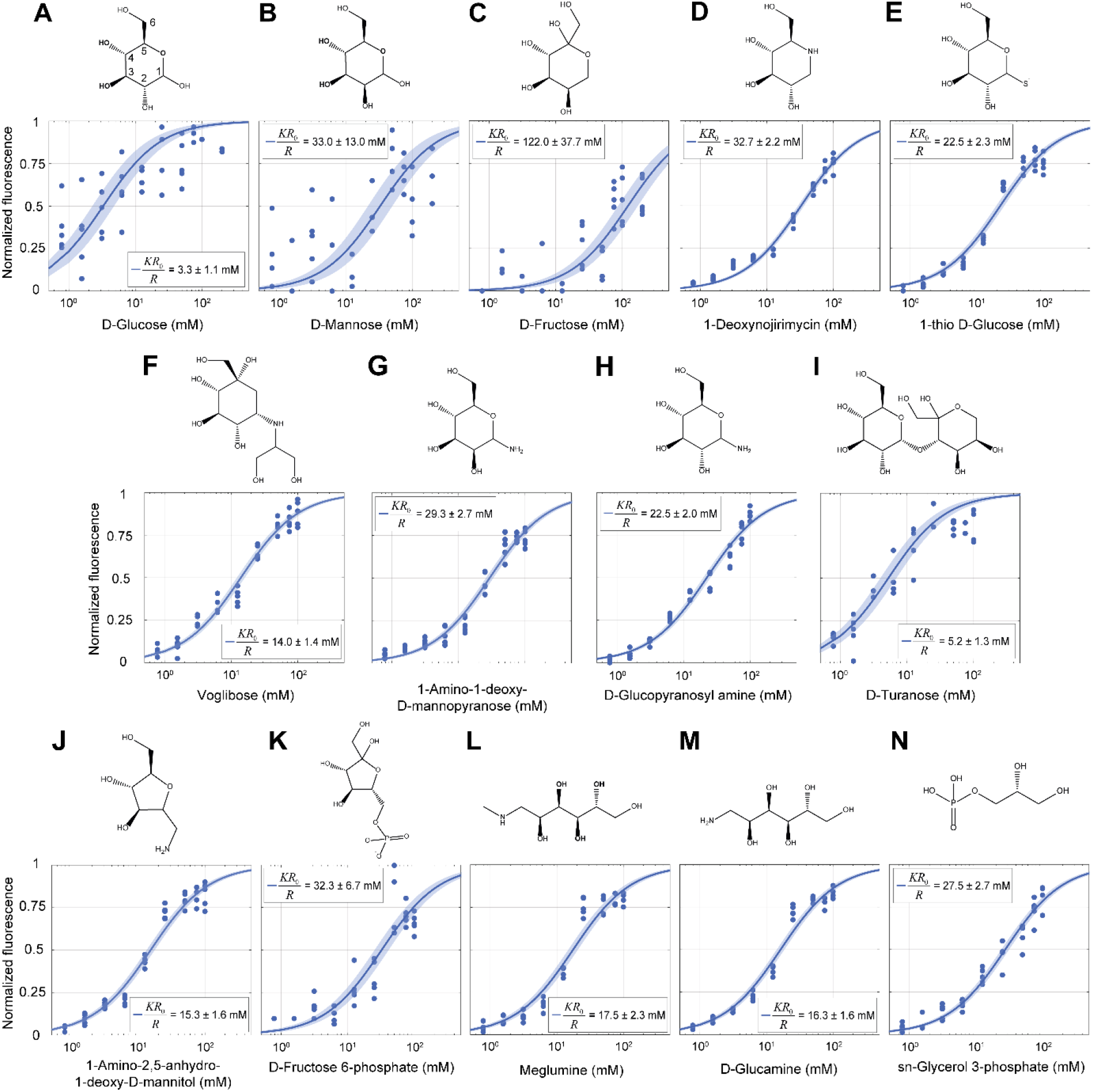
SweetTrac2′s steady-state response to sugar and sugar analogs transported by AtSWEET1. The carbons are numbered for D-glucose. Blue solid lines represent uniporter model fit as described in Park et al. 2022, and the shaded areas represent 95% confidence intervals. Equilibrium exchange constants are reported as estimated ± 95% confidence intervals (*n* = 5). All chemical structures are depicted in their most probable conformation in solution.

Given the high sequence identity of AtSWEET1 and AtSWEET2, we hypothesized that they would recognize similar substrates. Thus, the SweetTrac2 biosensor could help investigate whether the nine remaining hits identified in our previous work are bona fide substrates of AtSWEET1.

To this end, we first generated a yeast strain where AtSWEET1 is the sole hexose transporter on the plasma membrane. To achieve that, we integrated the *AtSWEET1* coding sequence into the genome of the EBY4000 strain, which lacks all endogenous hexose transporters. Subsequent expression of SweetTrac2 in the vacuole of these modified cells allowed us to test if any of the chemicals could be taken up via AtSWEET1.

SweetTrac2 showed increased fluorescence in response to 14 out of the 15 hits that were discovered in our previous work (Fig. 2), confirming that AtSWEET1 can mediate the cellular uptake of the majority of these chemicals (17). The only molecule that did not produce a fluorescence response was 1-deoxy-1-morpholino-D-fructose, suggesting that this modified sugar may be a competitive inhibitor of AtSWEET1 rather than a substrate or that AtSWEET2 may not be able to bind it.

Next, we measured the fluorescence response of SweetTrac2 to different concentrations of the 14 chemicals using our engineered strain. However, modifications were required to quantify the affinity of SweetTrac2 for D-glucose, D-fructose, and D-mannose, as the catabolism of these sugars produced variability in fluorescence measurements. We reasoned that higher amount of *AtSWEET1* template *DNA* would result in higher levels of protein and higher rates of sugar uptake, allowing cytosolic and extracellular concentrations to equilibrate faster and offsetting the consumption of these sugars by glycolysis. Accordingly, we overexpressed AtSWEET1 using a multicopy-number plasmid in EBY4000 instead of relying on the engineered strain where AtSWEET1 is integrated into the genome. These resulted in less variable fluorescence responses to different concentrations of the sugars (Fig.2A-C). The remaining 11 chemicals were tested our engineered strain (Fig.2D-N).

The affinity of SweetTrac2 for the different chemicals that produced a fluorescence response can be quantified using an equilibrium exchange constant (*KR*_0_/*R*), which we previously defined as the concentration of substrate that would saturate half of the biosensor at a steady state (14). SweetTrac2 displayed the highest affinity for D-glucose (*KR*_0_/*R =* 3.3±1.1 mM, Fig. 2A) and the lowest for D-fructose (*KR*_0_/*R =* 122.0±37.7 mM, Fig. 2C), intimating that D-glucose is the preferred substate of AtSWEET2.

We point out that the results for D-turanose need to be interpreted cautiously (Fig. 2I). Both EBY4000 cells expressing AtSWEET1 and cells transformed with an empty vector grow in the presence of D-turanose (Fig. S1). This suggests that there may be other transporters capable of mediating the uptake of D-turanose in the EBY4000 strain. Moreover, D-turanose is broken down by α-glucosidase in yeast cells into D-glucose and D-fructose (21). Hence, the steady-state response of SweetTrac2 to D-turanose is likely a composite response to the multiple sugars rather than only the disaccharide. This may explain why the *KR*_0_/*R* value for D-turanose is remarkably close to the value for D-glucose.

### Investigating the basis of substrate specificity using SweetTrac1

We were surprised to discover the differences in *KR*_0_/*R* between SweetTrac1 and SweetTrac2 (Table 1). Among the 14 chemicals that could bind SweetTrac2, we noticed that D-mannose, D-glucopyranosyl amine, D-turanose, 1-amino-2,5-anhydro-1-deoxy-D-mannitol, and *sn*-glycerol 3-phosphate (Fig. 2B,H-J,N) showed *KR*_0_/*R* values that were one order of magnitude lower than previously reported for SweetTrac1, while the *KR*_0_/*R* values for the other 9 chemicals were within the same order of magnitude for both biosensors (Table 1). We reasoned that the differences in *KR*_0_/*R* may be associated with differences between the substrate binding pocket of AtSWEET2 and AtSWEET1. Therefore, we decided to test this idea experimentally.

**Table 1.**
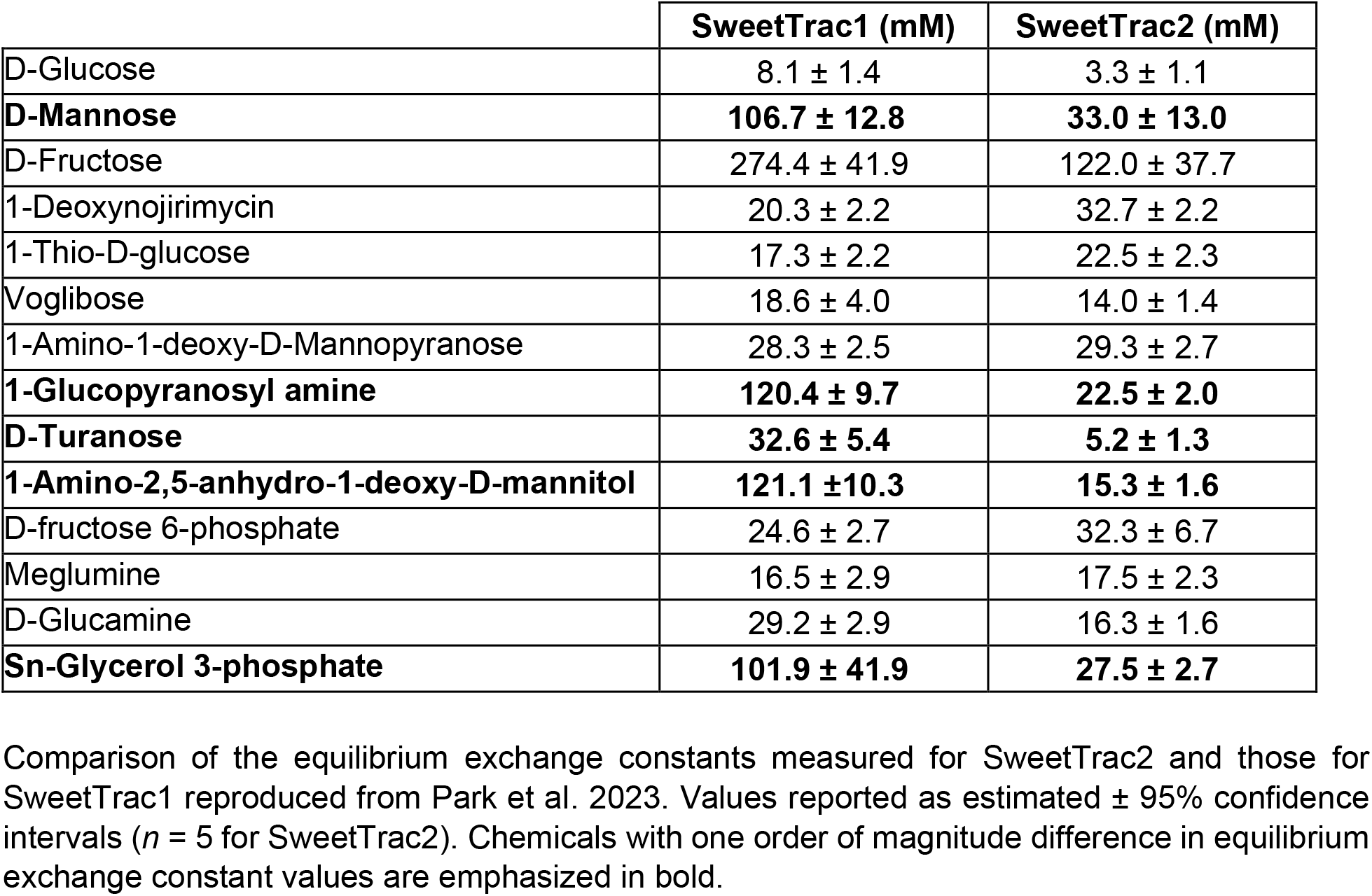
Steady-state response of SweetTrac1 and SweetTrac2 to different chemicals.

|  | **SweetTrac1 (mM)** | **SweetTrac2 (mM)** |
| --- | --- | --- |
| D-Glucose | 8.1 ± 1.4 | 3.3 ± 1.1 |
| **D-Mannose** | **106.7 ± 12.8** | **33.0 ± 13.0** |
| D-Fructose | 274.4 ± 41.9 | 122.0 ± 37.7 |
| 1-Deoxynojirimycin | 20.3 ± 2.2 | 32.7 ± 2.2 |
| 1-Thio-D-glucose | 17.3 ± 2.2 | 22.5 ± 2.3 |
| Voglibose | 18.6 ± 4.0 | 14.0 ± 1.4 |
| 1-Amino-1-deoxy-D-Mannopyranose | 28.3 ± 2.5 | 29.3 ± 2.7 |
| **1-Glucopyranosyl amine** | **120.4 ± 9.7** | **22.5 ± 2.0** |
| **D-Turanose** | **32.6 ± 5.4** | **5.2 ± 1.3** |
| **1-Amino-2,5-anhydro-1-deoxy-D-mannitol** | **121.1 ± 10.3** | **15.3 ± 1.6** |
| D-fructose 6-phosphate | 24.6 ± 2.7 | 32.3 ± 6.7 |
| Meglumine | 16.5 ± 2.9 | 17.5 ± 2.3 |
| D-Glucamine | 29.2 ± 2.9 | 16.3 ± 1.6 |
| **Sn-Glycerol 3-phosphate** | **101.9 ± 41.9** | **27.5 ± 2.7** |

Comparison of the equilibrium exchange constants measured for SweetTrac2 and those for SweetTrac1 reproduced from Park et al. 2023. Values reported as estimated ± 95% confidence intervals ( $n = 5$ for SweetTrac2). Chemicals with one order of magnitude difference in equilibrium exchange constant values are emphasized in bold.

Given our results and ample mutagenesis studies performed on AtSWEET1 (2), we propose that substrate recognition by SWEETs relies on a combination of specific and non-specific interactions. The specific interactions consist of hydrogen bonds formed between key hydroxyl groups in the substrates and conserved residues in the transporters, such as N73 and N192 in AtSWEET1 (2). We previously showed that mutating N73 and N192 abolished the fluorescence response of SweetTrac1 (14). Non-specific interactions are mediated by hydrophobic residues that determine the size and tortuosity of the binding pocket and may better explain the subtle differences in affinities between the two biosensors (Table 1). The role of the binding pocket size has been previously demonstrated for bacterial SemiSWEETs and the disaccharide transporter AtSWEET13 (5,22,23).

From molecular docking simulation results using the available crystal structure of the rice (*Oryza sativa*) SWEET2b (4) and D-glucose, D-fructose, and D-mannose, we selected 3 hydrophobic residues (V73, V76, and I193) in the binding pocket that are most likely to be directly involved in the interaction with the three sugars (Fig. 3). This corresponds to residues V69, I72, and V188 in AtSWEET1.

**Figure 3.**
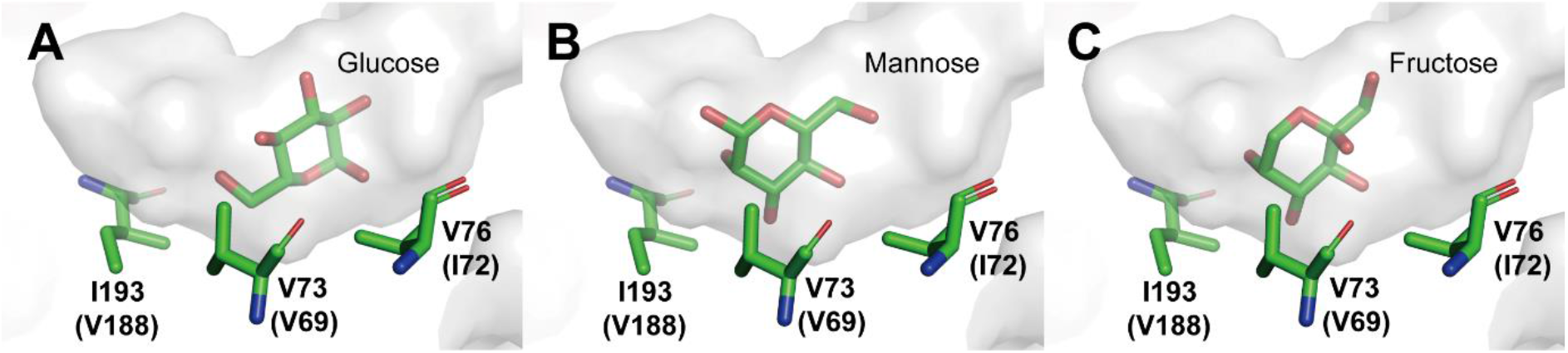
Molecular docking simulation results were performed on the OsSWEET2b with (A) D-glucose, (B) D-mannose, and (C) D-fructose. The substrate binding pocket is shown in light gray, and sugars and key amino acids in the binding pocket of OsSWEET2b are shown as sticks. Corresponding amino acids in AtSWEET1 are listed in parentheses. Mutations in V69, I72, and V188 altered the affinity of SweetTrac1 for D-glucose, D-mannose, and D-fructose. Image created with PyMOL Molecular Graphics System (Version 2.0 Schrödinger, LLC) from Protein Data Bank ID 5CTG.

Next, we performed alanine, isoleucine, and leucine substitutions in the V69, I72, and V188 residues of SweetTrac1 (numbering based on the sequence on AtSWEET1) (Fig. 1). Except for I72A, all other mutant biosensors correctly localized to the plasma membrane (Fig. S2) and showed a concentration-dependent change in fluorescence intensity (Fig. S3-S5).

Mutation of the less conserved V188 that made the binding pocket smaller (V188I and V188L) increased the biosensor’s affinity for D-glucose, D-mannose, and D-fructose (Table 2). In contrast, mutations that made the binding site bigger (V188A) had the opposite effect, albeit to different extents (Table 2).

**Table 2.** Steady-state response of SweetTrac1 mutants to three different sugars.

|  | D-Glucose<br>(mM) | D-Mannose<br>(mM) | D-Fructose<br>(mM) |
| --- | --- | --- | --- |
| SweetTrac1 | 8.1 ± 1.4 | 106.7 ± 12.8 | 274.4 ± 41.9 |
| V69A | 13.7 ± 2.2 | 558.5 ± 94.6 | 411.2 ± 122.0 |
| V69I | 1.5 ± 0.3 | 23.1 ± 5.4 | 72.3 ± 10.6 |
| V69L | 10.5 ± 1.7 | 311.7 ± 57.3 | 96.8 ± 17.3 |
| I72V | 6.2 ± 0.9 | 218.8 ± 30.3 | 334.4 ± 43.9 |
| I72L | 16.7 ± 3.4 | 173.0 ± 44.6 | 197.3 ± 36.3 |
| V188A | 20.8 ± 2.7 | 1977.0 ± 467.0 | 851.4 ± 255.4 |
| V188I | 7.0 ± 0.9 | 69.4 ± 13.1 | 173.6 ± 24.3 |
| V188L | 5.0 ± 0.7 | 29.1 ± 6.4 | 29.4 ± 5.1 |

Notably, some mutations of the more conserved V69 and I72 residues differentially affected the binding of D-glucose, D-mannose, and D-fructose. Specifically, V69L and I72L worsened the binding of D-glucose but facilitated that of D-fructose, while I72V had the opposite effect. All three mutations decreased the affinity of SweetTrac1 for D-mannose (Table 2). These results suggest that V69 and I72 may affect the shape or tortuosity of the binding pocket, which reduces steric hindrance for some substrates but increases it for others.

## Discussion

Due to their role in cellular energy and carbon storage, many studies have been done on vacuolar membrane sugar transporters, including the three vacuolar SWEETs (18,24-27). In this work, we generated SweetTrac2, a new vacuolar biosensor that reports the activity of AtSWEET2 *in vivo*. SweetTrac2 localized to the vacuolar membrane in yeast cells (Fig. 1A), mimicking the location of the natural transporter *in planta* (18). The ability to quantify the activity of transporters localized to intracellular membranes is especially significant since they cannot be studied in whole cells with radiotracer uptake or growth assays.

Since AtSWEET1 and AtSWEET2 share a considerable level of sequence identity, we used the same linkers and sfcpGFP in SweetTrac1 to convert AtSWEET2 into SweetTrac2. The success of this approach suggests that using plasma membrane transporters as proxies to convert intracellular membrane transporters into biosensors may be a viable method that bypasses the need to isolate organelles and reconstitute vesicles. However, we note that the approach may be limited to homologs with high sequence identity.

We explored the substrate specificity of AtSWEET2 using SweetTrac2 and discovered 14 sugars and sugar analogs capable of binding to the biosensor. Overall, it appears that the stereochemistry of the hydroxyl groups at the C3, C4, and C6 positions of the sugars (numbered according to D-glucose, Fig. 2A) are crucial for the recognition of substrates by AtSWEET2 as was previously suggested for AtSWEET1 (17). However, we noticed that the affinity of SweetTrac2 for some of these chemicals is distinct from SweetTrac1 (Table 1) and identified three hydrophobic residues in the binding pocket of AtSWEET1 that contribute to the differences.

Lastly, our work illustrates the use of biosensors to tune the specificity and selectivity of transporters and potentially other small molecule binding proteins. Mutagenesis analysis of the conserved V69 and I72 residues in the binding pocket of SweetTrac1 decreased affinity for D-glucose while increasing it for D-fructose, or vice versa. Future work combining several of these mutants or identifying new ones could help make D-fructose the preferred substrate of AtSWEET1 or even allow this transporter to recognize new substrates. The isolation of mutants that allow D-xylose transport would be particularly interesting, for example, by mutating the binding pocket of AtSWEET1 to resemble that of the D-xylose transporter AtSWEET7 (28). Such an engineered transporter could be beneficial for obtaining higher biofuel yields from hemicellulose fermentation (29).

## Methods

### DNA constructs

The *Arabidopsis SWEET2* (At3g14770) coding sequence was used as the basis for constructing the SweetTrac2 biosensor. SweetTrac2 contains the sequence of cpsfGFP, flanked by the left and right amino acid linkers DGQ and LTR, respectively, derived from the SweetTrac1 sensor (14). The cpsfGFP with linkers fragment was inserted after amino acid position 102 of AtSWEET2.

The AtSWEET1 (At1g24300) mutants were generated via QuickChange site-directed mutagenesis (Agilent) in pDONR-AtSWEET1 vectors. Sequences of SweetTrac2, AtSWEET1, and AtSWEET1 mutants were cloned into pDRf1-GW or p112A1NE using Gateway cloning (Invitrogen) for yeast expression. AtSWEET1 was cloned into pAG304GPD-ccdB (a gift from Susan Lindquist, Addgene plasmid # 14136) (30) using Gateway cloning (Invitrogen) for yeast genome integration.

### Yeast transformation and genome integration

*S. cerevisiae* strain EBY4000 (h*xt1-17D::loxP gal2D::loxP stl1D::loxP agt1D::loxP ydl247wD::loxP yjr160cD::loxP*)(31) and 23344c (*ura3*) were used in this study. Yeast was transformed using the LiAc/SS carrier DNA/PEG method and selected on solid synthetically determined, minimal medium (SD) supplemented with 0.67% w/v yeast nitrogen base without amino acids, 2% w/v maltose, and 0.19% w/v amino acids drop-out without uracil (Sigma Aldrich).

For yeast genome integration of AtSWEET1, pAG304GPD-AtSWEET1 was linearized using BstXI (NEB). 1 μg of linearized DNA was transformed into EBY4000 using the LiAc/SS carrier DNA/PEG method and selected on solid synthetically determined, minimal medium (SD) supplemented with 0.67% w/v yeast nitrogen base without amino acids, 2% w/v maltose, and 0.19% w/v amino acids drop-out without tryptophan (Sigma Aldrich).

### Fluorimetric analyses and parameter estimation

The excitation and emission spectra were measured in 23344c yeast cells expressing pDRf1-SweetTrac2 as described in Park et al. 2022.

Chemicals used in this study are listed in Table S1. The steady-state kinetic fluorescence response of SweetTrac2 was measured as previously described (17). To capture the steady-state fluorescence response for D-glucose, D-fructose, and D-mannose, EBY4000 cells co-transformed with p112A1NE-AtSWEET1 and pDRf1-SweetTrac2 were used. For all other molecules, EBY4000 cells with *AtSWEET1* integrated into the genome and transformed with pDRf1-SweetTrac2 were used. All chemicals were tested in SD medium to a final concentration of 0.78, 1.56, 3.13, 6.25, 12.50, 25, 50, 75, and 100 mM of sugars and a final OD_600_ of 0.5 and allowed to equilibrate for 6 minutes. We tested 5 replicates for each chemical. EBY4000 cells expressing the empty pDRf1 vector were used for background subtraction.

We noticed that 1-deoxynojirimycin, voglibose, 1-thio-D-glucose, meglumine, and D-glucamine had a negative effect on the baseline fluorescence when tested at high concentrations (around 10 mM) (Fig. 2D-K,L-M). This suggests that meglumine and D-glucamine may hinder cell metabolism in a similar manner to that previously demonstrated for 1-deoxynojirimycin, voglibose, and 1-thio-D-glucose (5,32). Subtracting the background during the analysis helps to minimize this hindrance from affecting our analysis, yet it assumes that each chemical has an equal effect on the cells that are expressing pDRf1-SweetTrac2 or empty vector.

### Yeast imaging conditions

A Zeiss LSM 700 Laser scanning confocal microscope with a 63x oil objective was used to visualize the localization of SweetTrac2 in yeast (Fig. 1C). The engineered EBY4000 strain expressing pDRf1-SweetTrac2 was used for imaging. The yeast culture was washed twice with deionized water and resuspended in 40 mM PBS with 200 mM glucose at pH 6.0. A 488 nm laser line was used to excite the fluorophore, and a 525/50 nm filter set was used to collect the emission (Carl Zeiss Microscopy). All images were processed with ImageJ.

### Molecular docking simulations

All ligand docking was performed on the rice OsSWEET2b (PDB ID 5CTG). The protein and ligands were converted to PDBQT format using AutoDockTools 1.5.7 (Dec 19, 2018) (33). The docking was performed with AutoDock Vina 1.1.2 (May 11, 2011) (34). All parameters were set to default values, and the search space was selected based on the substrate known binding site of the transporter. Results generated by the docking program were directly loaded into PyMOL Molecular Graphics System (Version 2.0 Schrödinger, LLC) and analyzed. All images were created with PyMOL.

## Acknowledgments

We are grateful to Taylor M. Chavez for assistance with experiments. This material is based upon work supported by the National Science Foundation under Grant No. 1942722 and the International Human Frontier Science Program Organization under Award No. RGY0077.

The authors declare that they have no conflicts of interest with the contents of this article.

