## Supplemental Information for "The Arabidopsis SWEET1 and SWEET2 uniporters recognize similar substrates despite differences in subcellular localization"

### Additional Methods

#### Yeast transformation

All constructs were transformed into *S. cerevisiae* strain EBY4000 (h*xt1-17D::loxP gal2D::loxP stl1D::loxP agt1D::loxP ydl247wD::loxP yjr160cD::loxP*) (31) using the lithium acetate method. Cells were plated on synthetically determined, minimal medium (SD) supplemented with 0.67% w/v yeast nitrogen base without amino acids, 2% w/v agar, 2% w/v maltose, and 0.19% w/v amino acids drop-out without uracil (Sigma Aldrich).

#### Growth assay

Yeast cells (strain EBY4000) expressing pDRf1-AtSWEET1 and cells transformed with the empty vector were used to test whether yeast can metabolize D-turanose (Fig. S1). 1.8 mL of cell suspension was added to 200 µl of 10X concentrated SD medium containing D-turanose as the only carbon source, resulting in a final concentration of 100 mM D-turanose and a cell $OD_{600}$ of 0.2. The cultures were stirred and incubated at 30°C for 8 hours before measuring their $OD_{600}$ using a Tecan Spark plate reader.

#### Yeast imaging conditions

Yeast cells (strain EBY4000) expressing SweetTrac1 mutants were visualized using a confocal microscope. Cells were washed twice with water and resuspended in 40 mM PBS with 100 mM D-glucose at pH 6.0 before imaging. A Zeiss LSD 700 laser scanning confocal microscope with a 63x oil, 1.4 NA objective was used. Fluorophore was excited using a 488 nm laser line, and emission was collected using a 525/50 nm filter set (Carl Zeiss Microscopy). All images were processed using ImageJ.

#### Fluorimetric analyses and parameter estimation

The steady-state kinetic fluorescence response of SweetTrac1 mutants was measured as described in Park et al. 2023 (17).

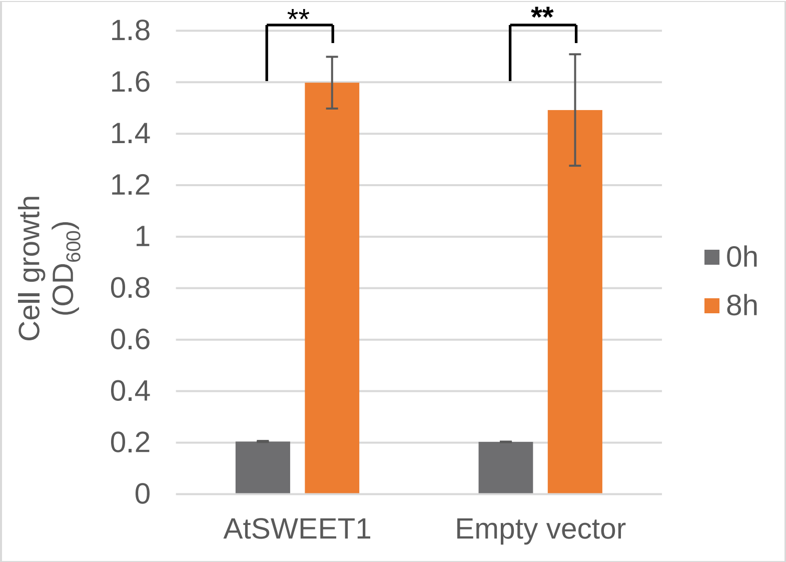


**Figure S1.** Transport of D-turanose by AtSWEET1 and subsequent growth in the yeast strain EBY4000. Cells were supplemented with 100 mM of D-turanose and incubated at 30°C for 8 hours. Optical density at 600 nm ($OD_{600}$) is reported as mean ± SD (*n* = 4). EBY4000 cells transformed with the empty vector served as the negative control. Significance according to student's t-test: ** $p<0.01$.

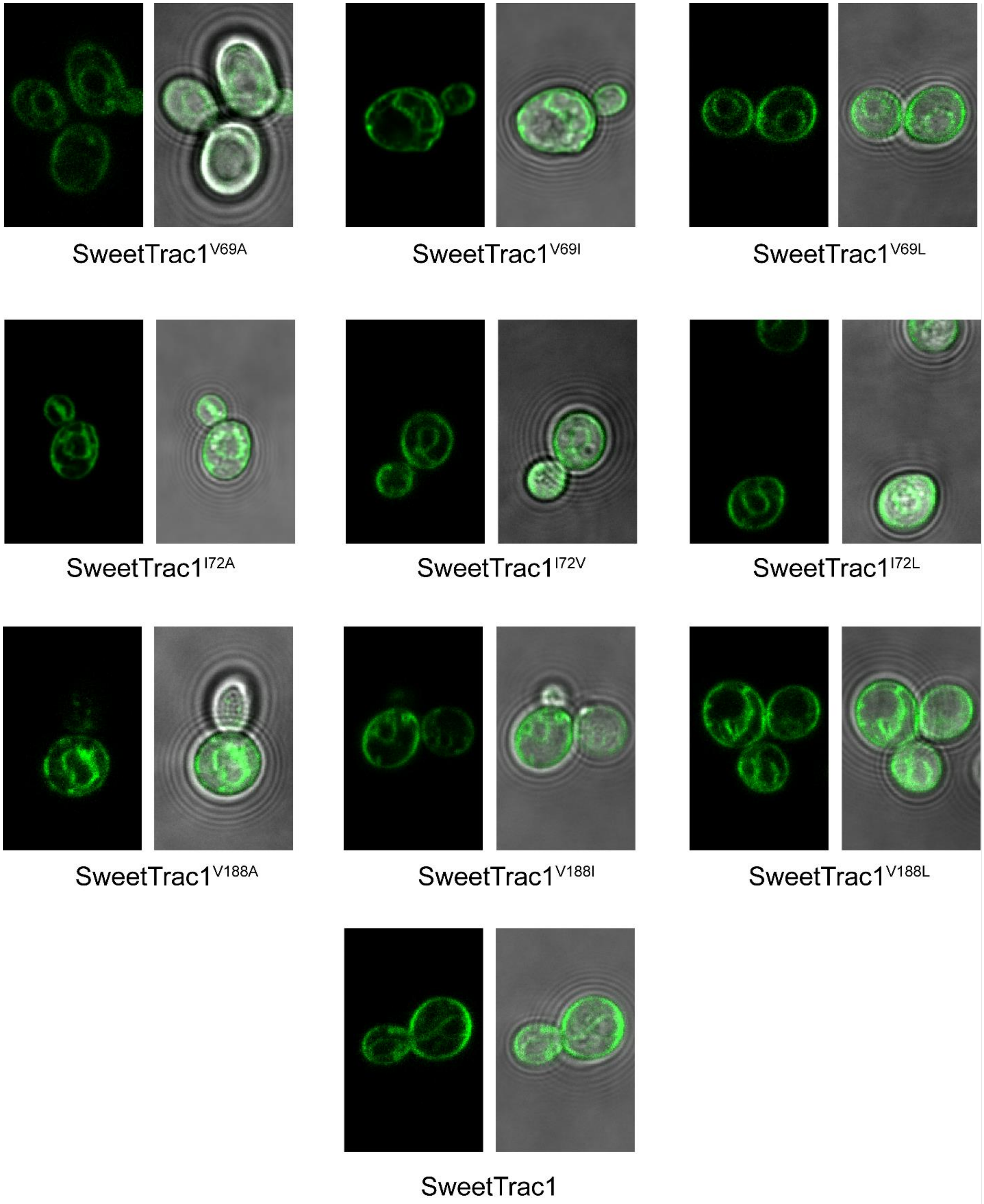


**Figure S2.** Localization of SweetTrac1 mutants in yeast. All mutant biosensors localized to the plasma membrane.

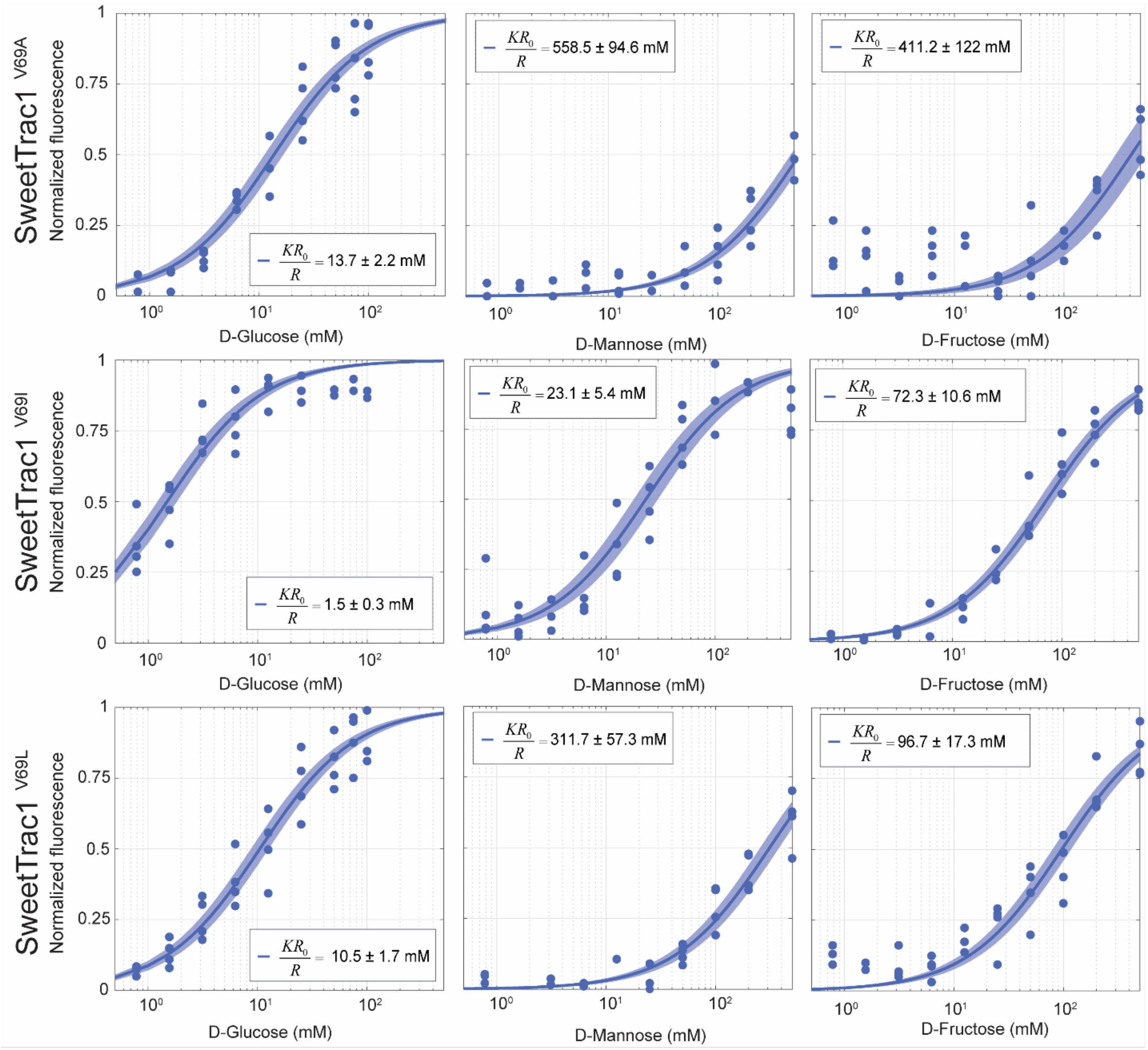


**Figure S3.** Steady-state kinetics of V69 SweetTrac1 mutants measured in microplate readers. Blue solid lines represent uniporter model fit, and the shaded areas represent 95 % confidence intervals. Equilibrium exchange constants are reported as estimate ± 95 % confidence intervals ($n$ = 4).

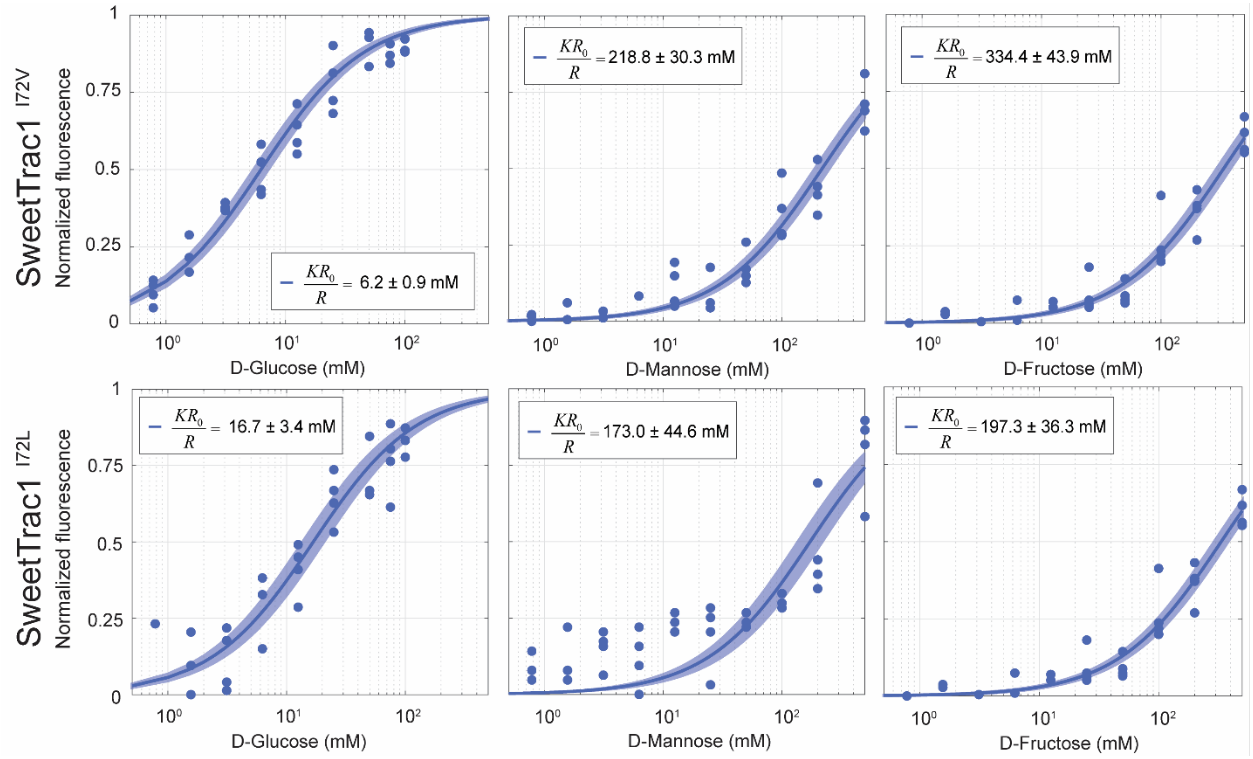


**Figure S4.** Steady-state kinetics of I72 SweetTrac1 mutants measured in microplate readers. I72A did not show a concentration-dependent fluorescence response. Blue solid lines represent uniporter model fit, and the shaded areas represent 95 % confidence intervals. Equilibrium exchange constants are reported as estimate ± 95 % confidence intervals (*n* = 4).

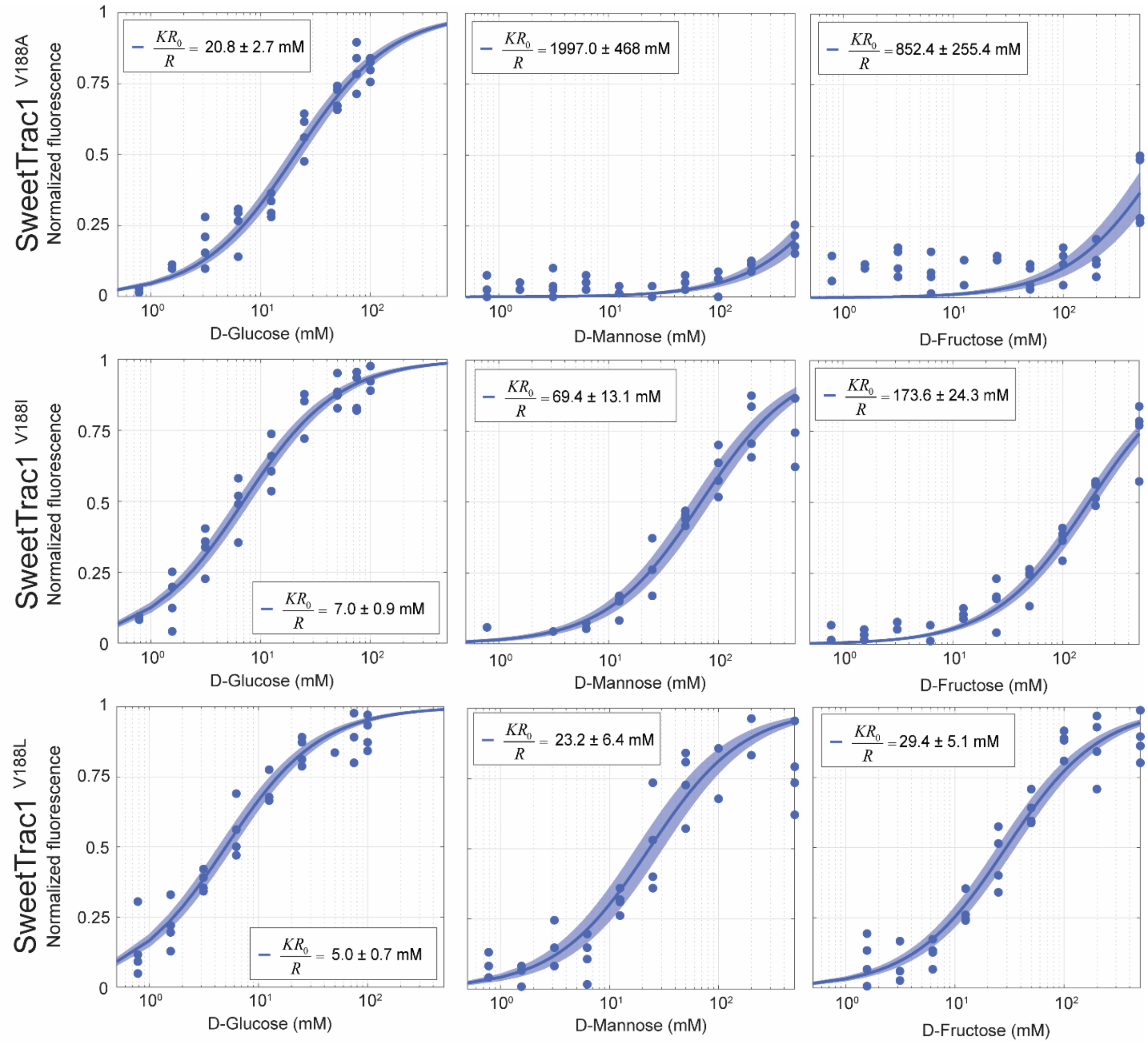


**Figure S5.** Steady-state kinetics of V188 SweetTrac1 mutants measured in microplate readers. Blue solid lines represent uniporter model fit, and the shaded areas represent 95 % confidence intervals. Equilibrium exchange constants are reported as estimate ± 95 % confidence intervals ($n$ = 4).

**Table S1. List of suppliers and catalog numbers for chemicals used**

| | Suppliers | Catalog # or SKU |
|---|---|---|
| D-Glucose | Sigma-Aldrich | G8270-1KG |
| D-Mannose | Sigma-Aldrich | M8574-500G |
| D-Fructose | Sigma-Aldrich | F0127-1KG |
| 1-Deoxynojirimycin | Biosynth International | MD05255 |
| 1-Thio-β-D-glucose sodium salt | Sigma-Aldrich | T6375-1G |
| Voglibose | Santa Cruz Biotechnology | sc-204384A |
| 1-Amino-1-deoxy-D-Mannopyranose | Santa Cruz Biotechnology | sc-364739A |
| β-D-Glucopyranosyl amine | Santa Cruz Biotechnology | sc-284974 |
| D-Turanose | Sigma-Aldrich | T2754-1G |
| 1-Amino-2,5-anhydro-1-deoxy-D-mannitol | Santa Cruz Biotechnology | sc-220459 |
| D-fructose 6-phosphate dipotassium salt | Sigma-Aldrich | F1502-1G |
| Meglumine | Sigma-Aldrich | M9179-100G |
| D-Glucamine | Biosynth International | MG06629 |
| sn-Glycerol 3-phosphate bis(cyclohexylammonium) salt | Sigma-Aldrich | G7886-1G |
